# Unsupervised discovery and predictive sensorimotor transformation of spider prey capture through active vibration sensing

**DOI:** 10.1101/2025.06.08.658484

**Authors:** Hsin-Yi Hung, Abel Corver, Andrew Gordus

**Affiliations:** Solomon H. Snyder Department of Neuroscience, Johns Hopkins University, Baltimore, MD; Department of Biology, Lund University, Lund, Sweden; Department of Biology, Johns Hopkins University, Baltimore, MD

## Abstract

Animals flexibly adjust posture and movement in response to vibrational sensory input to extract information from dynamic environments. While sensorimotor transformations have been extensively studied in visual and somatosensory systems, their structure remains poorly understood in substrate-borne vibration sensing. Here, we combine high-resolution web vibration recordings with fine-scale behavioral tracking in the orb-weaving spider *Uloborus diversus* to dissect the sensorimotor basis of prey capture.

Using unsupervised modeling, we identified discrete behavioral states that structure spider capture sequences, achieving over 83% classification accuracy. We then developed a predictive framework combining a linear-filtered generalized linear model (GLM) with a hidden Markov model (HMM) that robustly forecasts behavioral transitions across diverse prey vibration contexts. Notably, spiders exhibit context-dependent motor transitions—such as crouching and shaking—following decreases in prey vibrational power, consistent with active sensing behaviors that enhance signal detection. Furthermore, spiders reliably turn toward the web radius exhibiting the highest vibration amplitude during prey localization, demonstrating that amplitude alone predicts turning direction.

These findings reveal a structured, predictive sensorimotor transformation linking external vibration cues to internal behavioral states. Our results highlight general principles of active sensing and closed-loop control in non-visual invertebrate systems, with broader implications for sensorimotor integration across species.

## Introduction

Animals continuously acquire information from their environment through sensory systems to guide context-dependent behaviors such as foraging ^1,2^ and courtship^3,4^. However, sensory input is often noisy or transient, particularly during rapid or dynamic interactions such as prey capture. To overcome this, many species employ active sensing strategies^5,6^—coordinated motor actions that enhance the acquisition of behaviorally relevant stimuli. Active sensing has evolved across a wide range of sensory modalities. It is observed in vision^7,8^ (e.g., saccades in primates, head and body movements in flies), audition^9–11^ (e.g., echolocation in bats and whales), olfaction^12^ (e.g., sniffing in rodents), electrosensation^13,14^ (e.g., tail movements in electric fish), and mechanosensation^15–19^ (e.g., whisking in rodents or antennal scanning in insects).

A central theme in active sensing is the closed-loop nature of sensorimotor control: organisms do not merely react to sensory input but shape it through structured motor output. This continuous feedback loop improves signal detection, resolves sensory ambiguity, and enables flexible behavioral adaptation to dynamic environments. Despite its broad significance, the underlying structure and function of these sensorimotor loops remain incompletely understood, particularly in multimodal naturalistic contexts and vibration-based modalities such as substrate-borne signals.

The orb-weaving spider, *Uloborus diversus*, offers a powerful model to study vibration-based sensorimotor strategies. Unlike many animals that rely on vision or audition, spiders primarily sense their environment through substrate-borne vibrations. Their legs are equipped with highly sensitive slit sensilla—mechanoreceptors embedded at leg joints—which enable the detection of nanometer-scale displacements at frequencies up to 5,000 Hz^20,21^. Importantly, the spider web functions not only as a sensory substrate that transmits external vibratory cues but also as an actuator that is continuously reshaped by the spider’s own movements. This dual role makes the spider-web system an ideal substrate to dissect how animals actively modulate the sensory landscape and how those modulations, in turn, influence behavior. Spiders exhibit prey capture behavior that involves active movements in a naturalistic, context-dependent setting. Previous work ^22^ has described stereotyped movements—such as crouching, turning, and shaking—during prey capture in response to vibratory stimuli. However, these descriptions have largely remained qualitative, leaving open key questions about the functional role of these actions in shaping sensory input and how the resulting feedback influences subsequent motor decisions.

To address these questions, we investigated whether specific behaviors in the orb-weaving spider *U. diversus*—particularly crouching and shaking—amplify prey-induced vibratory signals, consistent with a role in active sensing. We first quantified the vibratory stimulus landscape spiders encounter using high-resolution web recordings. Next, we used unsupervised modeling to define a set of discrete behavioral states from leg kinematics, establishing a quantitative framework for decoding the spider’s behavioral repertoire. To investigate the sensorimotor relationship between stimulus and action, we built a predictive model linking sensory input to behavioral state transitions by combining Generalized Linear Models (GLM) with Hidden Markov Models (HMM). Spectral analyses revealed that crouching and shaking actively increase the vibratory power of subsequent prey-generated signals on the web. We further show that the structure of incoming vibratory input predicts future behavioral states, demonstrating a context-dependent, closed-loop dynamic. Finally, we explored how spiders localize prey through these interactions, shedding light on spatial computation and closed-loop control in a non-model organism.

Together, our findings reveal a dynamic, bidirectional sensorimotor loop in *U. diversus* that embodies core principles of active sensing, stimulus-guided behavior, and adaptive sampling. We present the first quantitative model linking substrate vibrations to discrete, predictive behavioral states in spiders during prey capture, offering a generalizable framework for understanding active sensing across diverse systems. These insights raise broader questions about the evolutionary convergence of sensorimotor strategies across diverse sensory modalities and species.

## Results

### *Drosophila melanogaster* and *Drosophila virilis* produce low-frequency vibration of 2-50 Hz on the web

To investigate the spider’s vibration-based sensorimotor transformation, we first defined the stimulus landscape that drives prey capture behavior. We developed a custom behavioral recording system that simultaneously captured web vibrations using a top-view high-speed camera (1,000 Hz) and spider behavior using a side-view camera (100 Hz) (Fig. 1a). The top-view recordings resolved frequency components up to 500 Hz over 8.734-second sessions, with a ring of white LEDs providing illumination to enhance silk contrast. Web geometry was annotated using a custom vision-based tracking algorithm (Fig. 1b), and a U-Net model^23^ trained on 91 manually labeled webs was used to predict web structure across additional recordings. Vibratory activity was quantified by measuring pixel intensity fluctuations along silk threads and applying Fast Fourier Transform (FFT) analysis, enabling non-contact, high-resolution mapping of the full web’s vibratory landscape with fine spatial and temporal precision.

**Figure 1.**
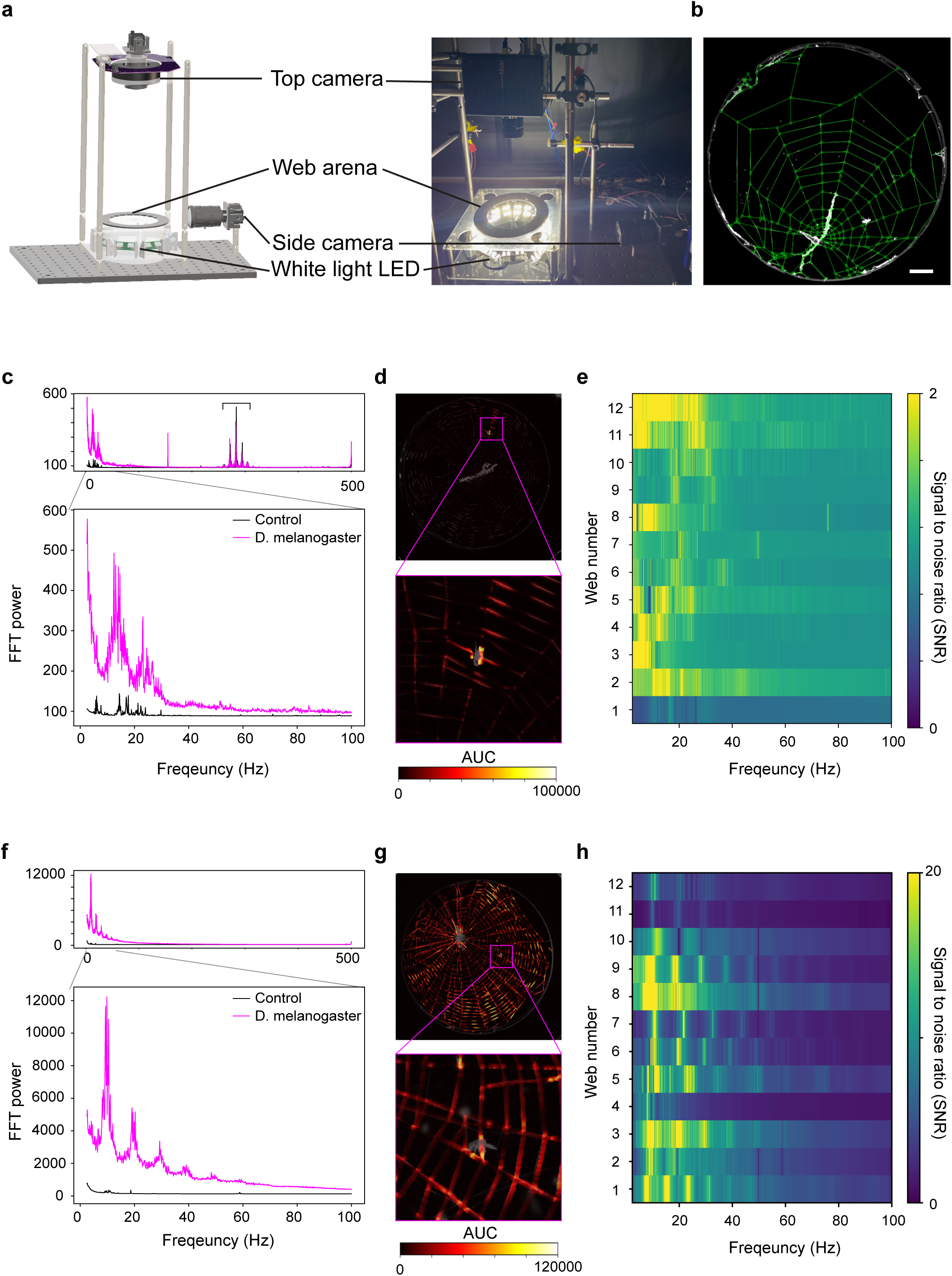
Web-borne vibrations induced by *Drosophila melanogaster* and *Uloborus diversus*. a, Schematic of the experimental setup for simultaneous recording of spider behavior and web perturbations. A ring of white LEDs illuminates the silk, while a high-speed top-view camera (1000 Hz) captures web vibrations and a side-view camera (100 Hz) records spider behavior. b, Example image showing annotated web architecture (green lines) used for vibration quantification. Scale bar = 1 cm. c, Representative fast Fourier Transform traces of vibrations induced by *D. melanogaster* (magenta) and in the absence of the fly (control, black), revealing broadband low-frequency activity with power concentrated between 2–50 Hz. Bracket region denotes narrowband noise frequencies. d, Area under the curve (AUC) map demonstrates that vibratory signals are spatially localized around the position of *D. melanogaster*. e, Signal-to-noise ratio (SNR), calculated as the ratio of Fourier power in the experimental versus control conditions, across 12 independent webs, confirms consistent *D. melanogaster*-induced 2-50 Hz vibration. f, *U. diversus* actively generates web vibrations during prey capture, producing resonance with a fundamental frequency at 10 Hz. g, AUC maps during spider-induced vibrations reveal widespread vibratory activity across the entire web. h, SNR analysis of spider-generated vibrations shows consistent harmonic peaks across 12 independent webs, highlighting the structured and resonant nature of self-generated signals.

We first recorded a baseline control for each web in the absence of any stimulus, followed by an experimental condition in which a *Drosophila melanogaster* was placed on the web. A representative example is shown in Fig. 1c, where *D. melanogaster* generates broad-band, low-frequency vibrations between 2-50 Hz. To further verify that these vibrations originated from the fly’s movements, we calculated the area under the Fourier transform curve (AUC) along each silk thread. We observed that AUC values peaked around the fly’s location, indicating that these signals primarily arise from *D. melanogaster* activity rather than background noise (Fig. 1d). We also computed the short-time Fourier transform (STFT) to examine the temporal patterns of prey signals, which exhibited irregular fluctuations (Supplementary Video 1). Next, we compared vibratory responses across 12 webs by computing the signal-to-noise ratio (SNR) as the ratio of the Fourier power spectrum in the experimental condition to that in the control (Fig. 1e). We found that *D. melanogaster* consistently produced broadband low-frequency signals in the 2-50 Hz range.

It is worth noting that in a subset of recordings, we detected narrowband noise peaks in both control and experimental conditions (bracket in Fig. 1c). Since these peaks were also present in the control and exhibited relatively low power (<600), we interpret them as likely artifacts, possibly arising from environmental or mechanical sources, rather than biologically relevant signals. Indeed, when we plotted the AUC map between 270–300 Hz, it revealed uniformly distributed power across the entire web (Supplementary Fig. 1a). The corresponding STFT further confirmed that this artifact persisted throughout the recording (Supplementary Fig. 1b), in contrast to the localized and irregular patterns characteristic of prey-induced vibrations.

To assess species-specific differences in vibratory signaling, we repeated the experiment using *Drosophila virilis*, a larger species of *Drosophila*^24^ (Supplementary Fig. 1c). Despite exhibiting a similar spectral range (Supplementary Fig. 1d–f), *D. virilis* generated substantially greater vibratory power on the web (Supplementary Fig. 1j). These findings indicate that species differences in web-borne signals arise primarily from vibratory intensity rather than frequency composition.

### Spiders actively produce harmonic vibrations on webs in response to *D. melanogaster*-generated cues

To determine whether spiders actively modulate web-borne vibrations in response to prey, we placed both *D. melanogaster* and *U. diversus* on the web. As a baseline, we first recorded vibratory activity with a stationary *U. diversus* in the absence of prey. Under these conditions, no significant vibrations were detected, indicating minimal background activity from the spider or ambient air currents (Fig. 1f). Upon the introduction of *D. melanogaster*, however, spiders actively generated vibrations by crouching or shaking webs, suggesting an active sensing mechanism triggered by the presence of prey. The AUC map reveals widespread vibration power across the entire web during active sensing (Fig. 1g, Supplementary Video 2). Interestingly, spider’s movements induce resonance on webs with 10-hertz fundamental frequency. These harmonic vibrations were consistently observed in 11 out of 12 recordings (Fig. 1h).

To probe whether this response depends on prey signal strength, we repeated the experiment with *D. virilis*, which generates higher-amplitude vibrations (Supplementary Fig. 1j). As predicted, spiders responded more rapidly to *D. virilis* than to *D. melanogaster* (Supplementary Fig. 1l). Notably, unlike the consistent harmonics observed with *D. melanogaster*, harmonic peaks at 25, 50, and 75 Hz were detected in only one of 12 trials with *D. virilis* (Supplementary Fig. 1g-i, k).

Together, these findings suggest that *U. diversus* produces harmonic vibrations in response to weak vibratory cues, potentially to enhance signal detectability via resonance. In contrast, when interacting with prey that already produces strong vibratory signals (*D. virilis*), the spider does not generate additional harmonics and reaches the prey faster.

### Unsupervised modeling reveals structured state transitions and timing modulation in spider prey capture across species

To observe how the spider dynamically vibrated the web, we tracked and quantified spider leg movement by using a side camera recording at 100 Hz (Figure 1a). We used DeepLabCut^25^ to track five joints on each of the two anterior and two posterior legs (20 joints total). Wavelet analysis^26^ was then applied to quantify the spiders’ limb movements (Fig. 2a). From the joint wavelet spectrum, 3 behavioral states were observed, which we named static state, crouching state, and high-frequency state. The static state was defined by low power in the spectrum due to very little limb movement (Supplementary Video 3). When the spider crouched, the joints moved in the 3-10 Hz frequency range. During high-frequency shaking states, the joint wavelet spectrum exhibited a prominent peak at 10 Hz—closely matching the resonance frequency identified in top-view recordings. This state could occur while the spider remained in one location on the web, or during walking or turning.

**Figure 2.**
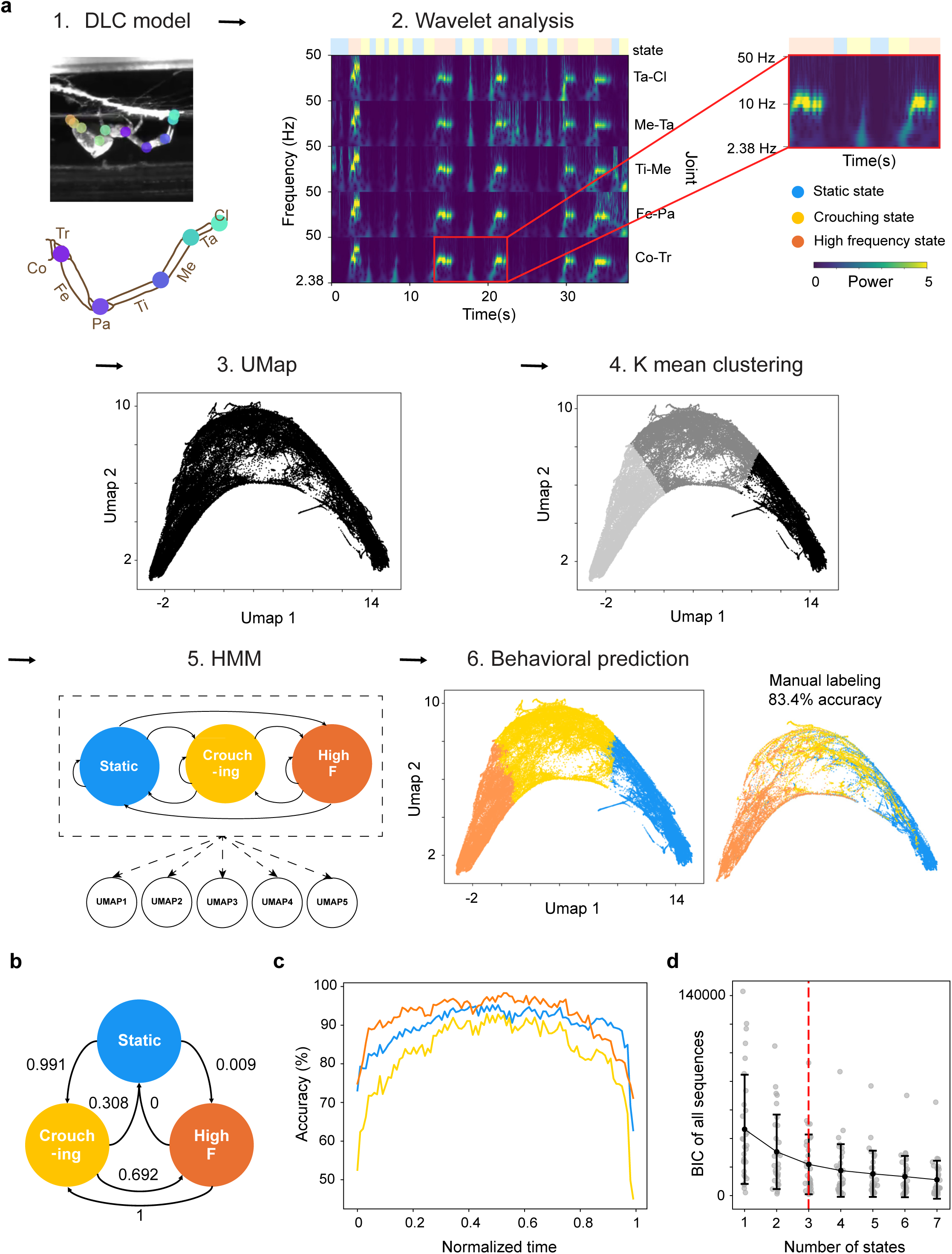
Unsupervised identification of spider prey capture dynamics during interactions with *D. melanogaster* using behavioral state modeling. a, Overview of the behavioral analysis pipeline. (1) Side-view recordings were analyzed with DeepLabCut to track 20 leg joints. Diagram denotes the joints tracked. (2) Wavelet spectrograms were computed from five joints on the left anterior leg (left), with a magnified view of the red-highlighted segment (right). Ethogram above the spectrogram denotes behavioral states. (3) UMAP was applied to reduce the joint dynamics to five dimensions across 30 recordings (UMAP dimensions 1 & 2 are shown). (4) Unsupervised K-means clustering in the UMAP space identified three distinct behavioral clusters. (5) A Hidden Markov Model (HMM) was then trained using the cluster-derived probability distribution to model the emission probabilities. (6) The HMM predicted behavioral states across all 30 videos, achieving 83.4% accuracy on 10 manually annotated sessions. Anatomical diagram: Co: Coxa, Tr: Trochanter, Fe: Femur, Pa: Patella, Ti: Tibia, Me: Metatarsus, Ta: Tarsus. b, HMM-inferred state transition probability matrix after removing self-transition. c, Time-normalized prediction accuracy of each state. Decreased accuracy is observed near state transitions, reflecting increased temporal uncertainty during behavioral switching. d, Bayesian Information Criterion (BIC) for HMMs with increasing numbers of hidden states. A sharp decrease from one to three states is observed, after which improvements plateau, supporting the selection of a three-state model. Dark grey circles represent individual sequences (30 videos). Data are presented as mean ± s.d. (filled circles and bars).

To automatically classify these states, we developed an unsupervised modeling pipeline combining dimensionality reduction, clustering, and Hidden Markov modeling (HMM) to predict a spider’s behavioral states based on joint wavelets. As the four legs exhibit similar wavelet patterns during prey capture (Supplementary Fig. 2a) and the spider’s primary motor output is driven by the anterior legs, we performed UMAP^27^ dimensionality reduction on five joints of the left anterior leg across 30 spiders to capture representative motor dynamics, yielding a five-dimensional embedding. Then, we applied unsupervised K-means clustering to the UMAP space, identifying three clusters to extract the probability distribution of three states. A Hidden Markov Model (HMM) was constructed using the five-dimensional UMAP embeddings as observables, with the emission probability distribution determined by K-means clustering^28^.

This unsupervised HMM enabled moment-to-moment behavior prediction, achieving 83.4% accuracy across 10 manually annotated trials out of 30 recordings with *D. melanogaster* interactions. Our HMM revealed a high probability of self-transitions within each behavioral state during prey-capture dynamics (Supplementary Fig. 2b). To better visualize the transitions between distinct states, we excluded self-transitions and normalized the remaining state-to-state transition probabilities (Fig. 2b). Notably, the model predicted all high-frequency states are followed by crouching states, likely reflecting biomechanical constraints that prevent abrupt cessation of movement. Overall, model performance declines during state transitions, but achieves 70–100% accuracy during non-transition periods (Fig. 2c).

To validate the number of behavioral states, we evaluated model performance using the Bayesian Information Criterion (BIC). BIC values dropped sharply between one and three states, followed by a plateau, indicating minimal improvement with additional states (Fig. 2d). We further quantified BIC changes relative to both the preceding state model and the best-fitting model per sequence (Supplementary Fig. 2c-d). In both cases, the greatest improvement occurred at three states, with marginal gains thereafter. Together, these findings support a three-state model underlying spider prey capture dynamics.

To test the generalizability of this three-state classification pipeline, we applied it to recordings of spiders interacting with a second prey species, *D. virilis*, which produces comparable vibratory frequencies (Supplementary Fig. 1j). Joint wavelet data from 12 *D. virilis* trials were projected onto the *D. melanogaster*-derived UMAP embedding (Supplementary Fig. 2e), and a new HMM was trained within this shared low-dimensional space. Using four manually labeled sequences for validation, the *D. virilis* model achieved 83.9% accuracy, comparable to that of the *D. melanogaster*-trained model. Importantly, transition dynamics were also qualitatively similar across prey types (Supplementary Fig. 2f), suggesting conserved underlying sensorimotor structure and supporting the robustness of the modeling framework.

Given that *U. diversus* responds more rapidly to *D. virilis* (Supplementary Fig. 1l), we hypothesized that this behavioral efficiency might arise from reduced engagement in crouching or shaking, or from shorter durations within each behavioral state. While the number of static, crouching, and shaking events did not differ significantly between prey types (Supplementary Fig. 2g), state durations were significantly shorter during *D. virilis* interactions (Supplementary Fig. 2h). In particular, crouching dwell times were reduced (Supplementary Fig. 2i), suggesting faster behavioral transitions and a larger decay constant. Together, these results demonstrate that *U. diversus* exhibits structured and stereotyped motor dynamics during prey capture that are robust across prey types, yet flexibly modulated in timing by sensory input intensity.

### Spiders’ crouching and shaking increase sensory stimulus power from *Drosophila melanogaster*

To uncover the functional role of crouching and high-frequency shaking during prey capture, we examined how these motor states influence vibratory signals from prey. As previously described, *D. melanogaster* produces relatively low-amplitude vibrations on the web (Supplementary Fig. 1j), and spider exhibits significantly longer state durations during interactions with this prey compared to *D. virilis* (Supplementary Fig. 2h-i). We hypothesized that crouching and high-frequency shaking behaviors may actively modulate prey signals to enhance sensory gain.

Because spider movement itself induces substantial web vibrations (Fig. 1f–h), we sought to isolate prey-induced signals from those driven by the spider. To achieve this, we extracted pixel intensity fluctuations from silk threads in a region of interest (ROI) centered on the *D. melanogaster* and normalized these values by fluctuations in a surrounding peripheral area (Fig. 3a). This normalization yielded a “pure” fly signal from top-view recordings, isolating prey-specific motion independent of spider-induced noise.

**Figure 3.**
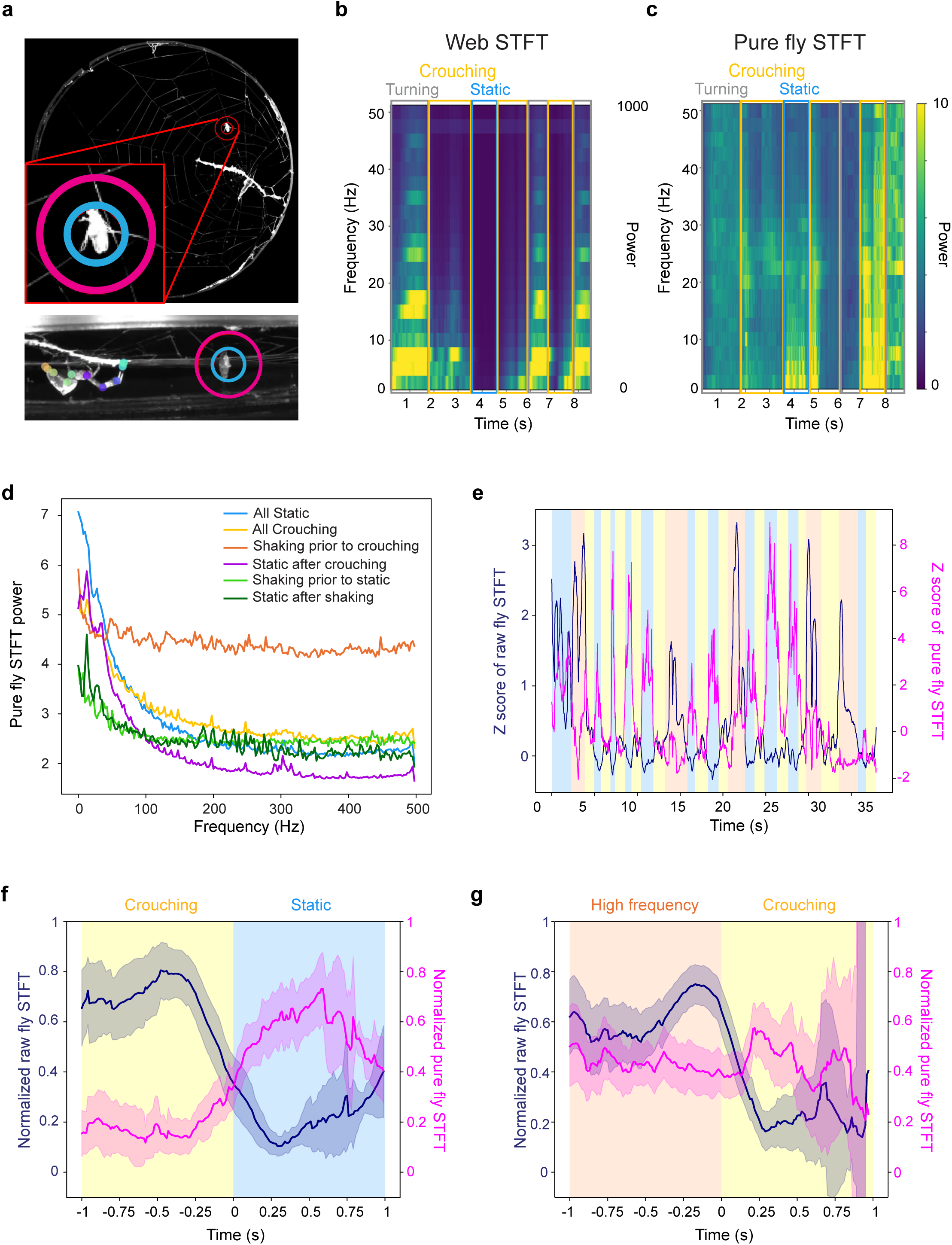
Spider crouching and high frequency movements increase sensory stimuli power from *Drosophila melanogaster*. a, Schematic illustrating regions of interest (ROIs) used to extract vibratory signals from the 1000 Hz top-view camera and behavioral states from the 100 Hz side-view camera. Area within the cyan circle was used to quantify the raw fly signal, which was normalized to signal between the magenta and cyan circles. b, Short-time Fourier Transform (STFT) power spectrum of the raw fly signal shows large amplitude fluctuations during spider movement, reflecting spider-induced web motion. c, STFT spectrum of the pure fly signal from the same recording as in b shows minimal power during spider turning, indicating passive fly movement. By contrast, crouching and static states are associated with elevated spectral power, suggesting active fly movement. d, Average short-time Fourier transform (STFT) power of pure fly signal across 12 top-view recordings reveals distinct spectral profiles across behavioral states. e, Example trace showing average raw and pure fly signal STFT power (0–30 Hz) from extended side-view recordings. Pure fly signal power increases during static periods following crouching. f, Quantification across 16 side-view recordings confirms consistent enhancement of low-frequency (0–30 Hz) normalized pure fly signal power during static states following crouching. Shaded areas represent 95% confidence intervals. g, Similar enhancement is observed during transitions from shaking to crouching, as predicted by the HMM. Shaded areas represent 95% confidence intervals.

Short time Fourier Transform (STFT) was then applied to the pure fly signal to investigate temporal dynamics of fly movements. We manually annotated spider’s behavioral states in the top-view data because state predictions from the HMM were based on lower frame rate side-view recordings at 100 Hz. During large spider movements in high-frequency state, such as turning, the raw fly signal displayed large power (Fig. 3b, Supplementary video 4), whereas pure fly signal exhibited low power (Fig. 3c), indicating that the fly moved passively with the vibrating web. By contrast, crouching behavior induced greater pure fly signal power compared to high-frequency state. Strikingly, upon cessation of crouching, fly power further increased—exceeding levels observed during crouching—suggesting that the crouching behavior itself may trigger fly movement and enhance the resulting sensory signal from *Drosophila melanogaster*.

To quantify these effects across multiple *D. melanogaster* trials, we averaged STFT power over time from 12 top-view video recordings. In the pre-capture static state, fly-induced vibrations were predominantly low-frequency (<30 Hz; Fig. 3d, cyan). When the spider was crouching, the pure fly signal showed a spectral peak at 12 Hz (Fig. 3d, yellow). Notably, power in the subsequent static state increased further, peaking at 12 Hz and 33 Hz (Fig. 3d, purple). Shaking behavior typically elicited broadband fly signals spanning 0–500 Hz, with shaking preceding crouching (Fig. 3d, orange) exhibiting higher power than shaking followed by a return to static (Fig. 3d, light green). Static states following crouching (purple) or shaking (dark green) consistently exhibited increased vibratory power relative to the prior state, suggesting that these behaviors enhance transmission of fly-induced signals across the web.

To examine whether sensory gain from pure fly signals in static states consistently increased following crouching throughout the entire prey capture sequence, we analyzed behavioral state transitions predicted by the HMM using extended-duration side-view recordings. We applied the same ROI-based analysis to quantify fly signals; however, due to limited visibility of silk threads in the side view, we defined the normalized fly signal directly from pixel intensity of the fly itself, rather than silk fluctuations.

By averaging STFT power within the 0–30 Hz frequency band, we observed a robust increase in pure fly signal during static states that followed spider crouching (Fig. 3e–f, Supplementary video 5). Although the HMM did not predict any transitions from shaking to static states, we found that high-frequency shaking movements were consistently followed by elevated sensory signals during the subsequent crouching state (Fig. 3g), further supporting the role of both crouching and shaking in amplifying prey-derived vibratory input.

### Fly-induced sensory cues predict spider behavioral states

If spider behavior functions to enhance sensory gain, we next asked whether prey-induced sensory cues could, in turn, predict the spider’s behavioral responses. To test this hypothesis, we developed a hybrid sensorimotor model combining generalized linear models (GLMs) with hidden Markov models (HMMs) (Fig. 4a), allowing us to capture both stimulus-dependent dynamics and temporal structure in behavior.

**Fig. 4.**
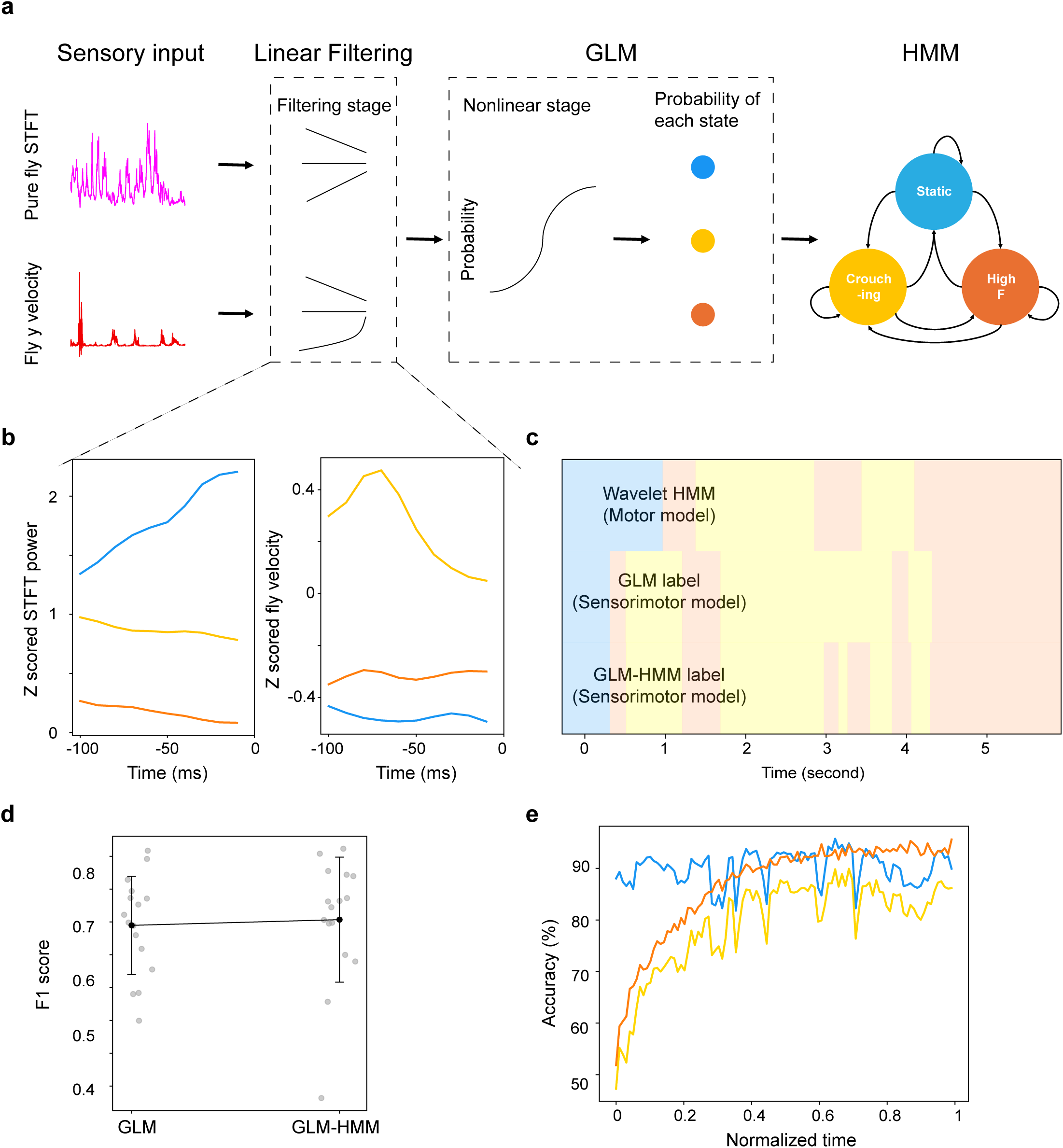
A sensorimotor model predicts spider behavioral states from prey-induced vibratory cues. a, Schematic of the GLM-HMM framework combining generalized linear models (GLMs) with hidden Markov models (HMMs) to predict spider behavior based on *D. melanogaster*-derived sensory input. The model incorporates two features extracted from Drosophila melanogaster: pure fly STFT power (magenta) and vertical fly velocity (red). To filter out noise in raw data, we estimate linear filters by using behavior- trigger average (BTAs). Inner product of raw sensory data with linear filters are passed through a nonlinearity logistic regression, providing the probability of each state on a moment-to-moment basis. Finally, an HMM is applied to incorporate the history of movement. b, Behavior-triggered averages (BTAs) showing sensory input aligned to the onset of spider behavioral states (cyan: static, straw: crouching, and orange: high-frequency movement). Static state transitions are preceded by an increase in fly vibration amplitude, while crouching and high-frequency movements are preceded by decreases in vibratory power. Spider’s crouching were preceded by reductions in fly vertical velocity. c, Example behavioral state sequences predicted by the wavelet HMM, GLM, and GLM-HMM models. The GLM-HMM produced more temporally coherent predictions than the GLM. d, F1 score comparison between GLM and GLM-HMM models across datasets (Mann–Whitney U test, p = 0.5591; filled circles denote mean ± s.d.), showing no significant difference in predictive accuracy. e, Time-normalized prediction accuracy of the GLM-HMM versus the wavelet-based HMM across 16 video recordings. The GLM-HMM achieved 75.3% accuracy while maintaining interpretability of stimulus–behavior relationships.

We analyzed 16 side-view recordings of spider–prey interactions and trained a multinomial GLM to classify spider behavior into three discrete states—static, crouching, and high-frequency movement— based on two sensory features from *Drosophila melanogaster*: the pure fly STFT power and fly vertical velocity. To examine how sensory input shapes behavioral transitions, we estimated linear filters — behavior-triggered averages (BTAs)—by aligning stimulus features over a 100-ms window preceding the onset of each behavioral state. Importantly, these linear filters revealed that fly vibratory power increased prior to transitions into the static state but decreased before the onset of crouching and high-frequency movements (Fig. 4b). Additionally, transitions into crouching were preceded by reductions in fly vertical velocity. Together, these results suggest that specific changes in prey-induced signals influence the probability of entering distinct behavioral states. More generally, spiders tend to pause their movements when the fly is moving, and crouch when the fly decreases its movement on the web.

We then used the filtered sensory signals—obtained by taking the inner product of the raw input and BTA filters—as input to a multinomial logistic function to predict behavioral state. Using 25% held-out data and 250 bootstrap iterations (n = 1000), the two-feature GLM significantly outperformed models using only STFT power (p < 10^-307^) or substituting horizontal for vertical velocity (p < 10^-307^), and performed comparably to the three-feature model (p = 0.683; Supplementary Fig. 3a). A 100-ms BTA window yielded optimal classification accuracy, with shorter (50 ms) and longer (250 ms, 500 ms) durations showing no improvement (Supplementary Fig. 3b). Also, filters estimated with alternative window lengths showed similar trends to those obtained with the 100-ms window (Supplementary Fig. 3c), indicating robustness of the derived sensory-behavior mappings.

Although the GLM successfully captured stimulus–response relationships, it lacked a mechanism for modeling temporal continuity between behavioral states. To address this limitation, we extended the model by incorporating a hidden Markov model (HMM), which jointly inferred both stimulus-driven behavioral outputs and state transitions. While the GLM and GLM-HMM exhibited similar prediction accuracies (Fig. 4c–d), the GLM-HMM yielded smoother behavioral sequences by integrating both instantaneous sensory inputs and state transition dynamics. The resulting GLM-HMM achieved 75.3% prediction accuracy across the 16 datasets compared to a wavelet-based HMM (Fig. 4e), providing interpretable insights into how specific vibratory features of prey stimuli govern behavioral state transitions in the spider.

### Spatial structure of vibratory input predicts orienting behavior

Finally, we examined whether spatial patterns of vibratory input guide the spider’s orienting behavior. Using top-view recordings, we quantified vibratory signals by measuring pixel intensity changes along individual radial threads preceding each turning event (Fig. 5a, Supplementary video 6), assessing whether localized variations in signal amplitude across the spider’s peripheral sensory field could predict turning direction on a trial-by-trial basis.

**Figure 5.**
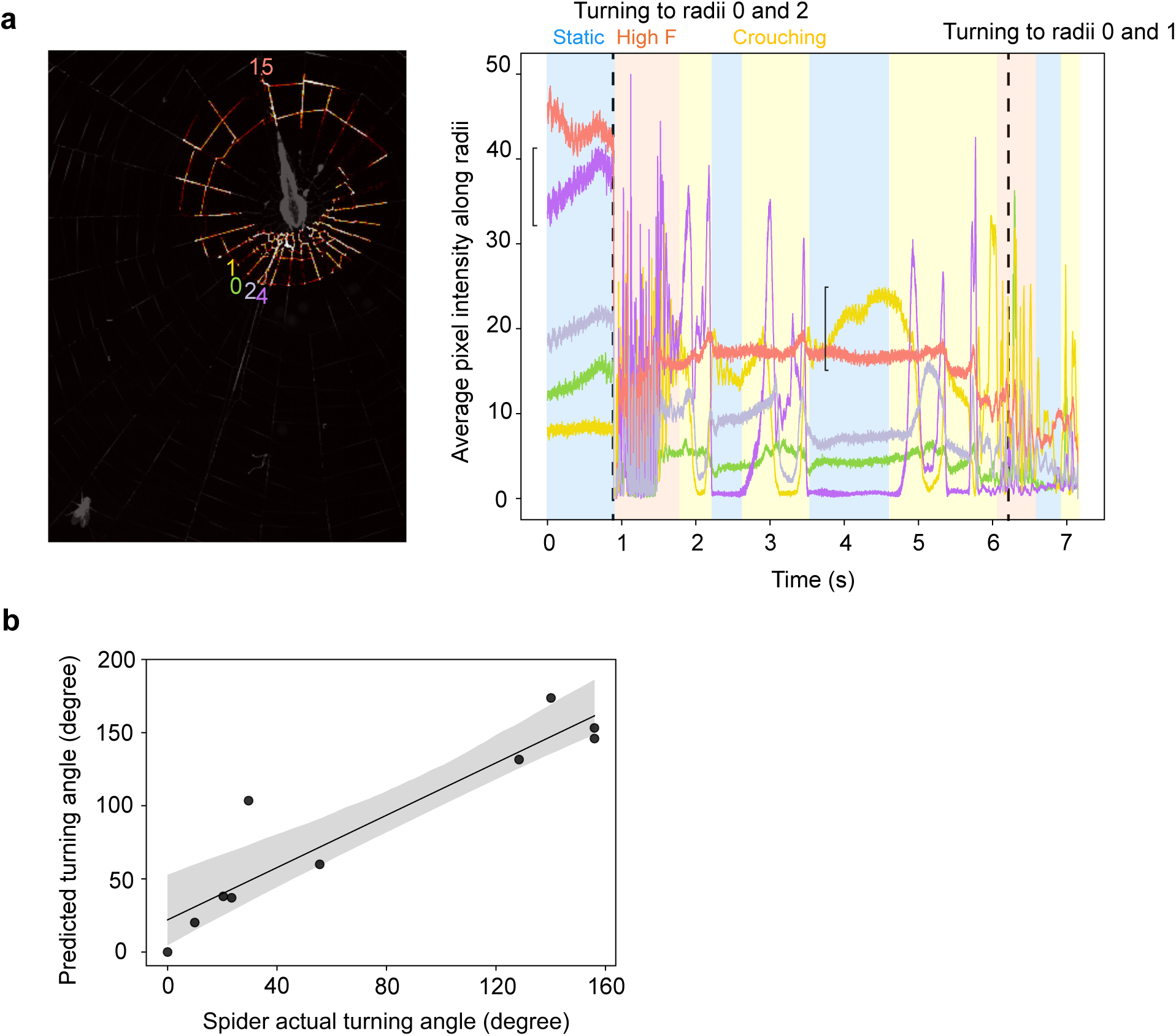
Spatially localized vibratory cues guide spider orienting behavior. a, Schematic of analysis pipeline using top-view recordings to extract vibratory signals from individual radial threads prior to spider turning events (left). Pixel intensity changes were quantified to estimate vibratory input along each thread within the spider’s peripheral sensory field (right). Bracket indicates largest amplitude change in static state prior to turning. Ethogram denotes spider behavioral states. b, Across trials, the spider consistently turned toward the radial thread with the highest vibratory amplitude (p =1.027 × 10^-4^, r = 0.9289, slope = 0.8943). A shaded area around the regression line represents the 95% confidence interval.

For each turning event, we identified the radial thread exhibiting the greatest vibratory amplitude in static state prior to movement onset and compared its spatial position to the spider’s turning angle. This analysis revealed a consistent correspondence: the location of peak vibration reliably predicted the direction of turning (Fig. 5b). In nearly all cases, the spider turned toward the radial thread carrying the strongest vibratory signal.

These results indicate that spatially localized vibratory cues serve as directional guides during prey capture, suggesting that spiders extract positional information from the vibratory landscape of the web. This spatial encoding of sensory input provides additional evidence for a dynamic sensorimotor loop, in which environmental signals not only elicit behavior but are actively interpreted through structured motor responses. Our findings underscore the importance of peripheral sensory dynamics in guiding closed-loop prey localization.

## Discussion

Our study reveals that spiders implement predictive sensorimotor strategies during prey capture, relying on active sensing of web vibrations. By employing an unsupervised modeling approach, we were able to objectively identify discrete behavioral states within a continuous, wavelet-based behavioral space, achieving an impressive 83.4% accuracy when compared with human annotations. This level of fine-scale, 10-millisecond resolution behavior is typically prone to biases introduced by subjective human labeling. While previous studies have proposed data-driven methods to identify behavioral motifs in animals— such as stereotyped movements in flies^26^ and web-making behavior in spiders^29^, and hunting sequences in zebrafish^30^ —we extend these efforts by investigating a finer-timescale, continuous, vibration-evoked active sensing behavior. In addition, wavelet transforms captures temporal dynamics across multiple frequencies, offering a richer and more comprehensive representation of movement variability than postural space.

Beyond behavioral identification, we introduced a novel sensorimotor transformation model that combines a Linear Filter Generalized Linear Model (GLM) with a Hidden Markov Model (HMM), enabling us to predict moment-to-moment behavior based solely on sensory input. This model demonstrated that features of fly-induced web-borne vibrations and fly velocities alone are sufficient to predict spider behavior, suggesting a strong coupling between environmental feedback and motor control. Notably, we simplified the elegant GLM-HMM framework developed by Calhoun et al.^31^ by estimating filters using a linear behavior-triggered average, along with an uncoupled GLM-HMM framework. Our linear filters show that spiders initiate crouching and shaking behavior when the vibrational power from prey, *D. melanogaster*, decreases. This approach offers several advantages, including simplicity, computational efficiency, and ease of biological interpretation. Importantly, unlike more complex methods, this non-parametric approach avoids the issue of getting stuck in local optima, making it a robust and reliable solution.

This dual-framework approach—first uncovering the structure of behavior, then linking it to sensory input—reveals that spiders encode and respond to vibratory cues in a temporally structured and context-dependent manner. We further demonstrate that spider’s crouching and shaking enhance sensory gain during the subsequent static phase, suggesting an active sampling strategy during prey capture. Our findings highlight a dynamic interplay between sensing and behavior, where spiders modulate their own movements to better interpret complex, fluctuating vibrational inputs. Future work may explore how spider sensory neurons encode different vibrational frequencies and amplitudes, and how these signals modulate downstream behavioral responses.

A critical aspect of the spider’s sensory system lies in its ability to define web-borne prey signals. Previous studies have shown that prey signals typically span a broad frequency spectrum from 5 to 1000 Hz. However, these investigations have primarily focused on airborne vibrations ^20,32^ or vibrations transmitted through rigid substrates, such as banana plants ^33^. In contrast, entrapped insects in spider webs have been reported to produce low-frequency vibrations, typically in the range of 5–50 Hz ^20,34^. Most of this research has focused on vertical orb webs, such as those built by *Nephila clavipes,* by using laser Doppler vibometry —a technique that measures vibration at a single point. In our study, we use high-speed camera recording at 1000Hz with vision-based technique to measure both temporal and spatial vibration profile of entire web. Our results show that prey captured in the horizontal orb web of *U. diversus* also generated vibrations in the 2–50 Hz range. Importantly, by leveraging the high spatiotemporal resolution of our method, we found that radii with the greatest vibration amplitude is sufficient to predict spider’s orientation during prey localization. This finding extends our understanding of web-borne prey signals from the more commonly studied vertical orb webs to horizontal orb webs, and shed light on the underlying mechanisms of spider prey localization.

Our findings offer valuable biological insights into the spider’s active vibration sensing during prey capture, while also providing computational models for behavioral discovery and sensorimotor transformations. These results draw parallels to active sensing strategies observed in animals such as bats^35,36^, rodents^37^, and weakly electric fish ^38^. They highlight the broader principle that robust behavioral sequences can be understood and predicted through an analysis of the sensorimotor loop, even in non-model invertebrate systems. Overall, this work advances our understanding of the complex, adaptive nature of behavior in response to sensory inputs, with implications for both fundamental science and the development of robotics and artificial intelligence.

## Methods

### Animals

*Uloborus diversus* were housed in an on-campus greenhouse at Johns Hopkins University. All animals were transferred to custom indoor habitats and kept on a 12 hr:12 hr light-dark cycle (15-30°C, 50%–70% RH) at least a week before being used for behavioral experiments. Spiders were fed *Drosophila melanogaster* or *Drosophila virilis* once a week. Only adult females were used in this study as adult males do not build orb webs.

### Behavioral assay

Adult females were placed in an arena with a 10 cm × 10 cm perimeter, coated with paper at the edges to encourage web-building. We used a ring of white light LED (Lite-On Inc., LTW-2S3D8) to illuminate spider silk. To increase imaging contrast, a high-absorption background material was placed below the behavioral arena (Acktar, Spectral Black Foil, SB-20x030-1-010). A Photron FASTCAM Mini UX100 high speed camera was set up on the top with a 16 mm fixed-focal length lens (Edmund Optics, #85-865) to record web vibration at 1000 Hz (1280 × 1024 pixel resolution). A side camera (BFS-U3- 16S2M-CS USB 3.1 Blackfly, 1440 × 1080 pixel resolution) with a 12 mm fixed-focal length lens (Edmund Optics, #86-570) simultaneously recorded spider’s movements at 100 Hz. PFV (ver.3610) and SpinView (1.27.0.48) software were used to store the recordings in AVI format.

Each web had two recordings: one control and one experimental. In the *Drosophila* vibration experiment, the control condition included an empty web, allowing us to measure baseline vibrations caused by air and terrestrial vibrations. The experimental condition included placing either *Drosophila melanogaster* or *Drosophila virilis* on the web to measure vibrations generated by prey movement. In the spider prey capture experiment, the control condition was of a live *Uloborus diversus* on the web without any perturbation. In the experimental condition, either *D. melanogaster* or *D. virilis* was introduced onto the web in the presence of the spider to record its behavioral response.

### Web annotation

We manually annotate 91 spider webs with previously developed an in-house web-tracking Graphical User Interface (GUI)^29^. 80% of the web data was used for training, 15% for validation, and 5% for testing. For each web, we computationally deleted parts of spider web and repeated this process 30 times, helping us to expand our dataset from 91 webs to 2821 webs (= 91 × 31). We train a U-Net^26^ to predict web structure with loss equal to 0.0475 and accuracy equal to 0.9821.

### Web vibration analysis

To extract the frequency profile of the web, Fast Fourier transform (FFT) analysis was applied on pixel intensities along silk lines. The FFT, of a time series with length *N*, was defined as

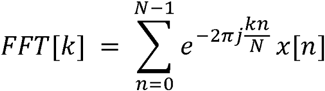

where *x*[*n*] is the discrete-time signal and *k* is the frequency bin index. The mean of FFT along all silk lines was computed to investigate whole web vibration.

To compare vibration patterns across all webs, we used signal-to-noise ratio (SNR) defined as

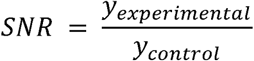

We also performed short-time Fourier Transform to investigate the temporal profile of web vibration. It was defined as

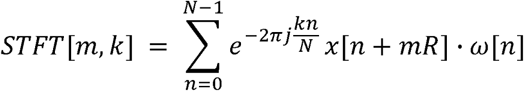

where *w*[*n*] is the window function of length 400 samples, *m* is the frame index, and R=20 is the hop size (i.e., the step between adjacent frames).

### Prey regions of interest (ROIs) analysis

To extract prey vibration on the web, we selected regions of interest around *Drosophila*. The prey’s peripheral field radius was twice the radius of the ROI, excluding the central ROI. Both FFT and STFT along silk lines were calculated within ROIs and peripheral field for top camera recordings. For the side camera, FFT and STFT were computed based on the pixel intensities of the fly and silk within the ROIs and the peripheral field, respectively.

Normalized STFT prey signal was defined as STFT within ROIs normalized by prey peripheral STFT.

### *U. diversus’* joint wavelet analysis

We used DeepLabCut^25^ to track 20 joints on spider’s anterior and posterior legs: body-coxa, coxa-femur, femur-tibia, and tibia-metatarsus joints, as well as the tip of the tarsus. Since spiders primarily move along the horizontal plane, all horizontal coordinates were centered by subtracting each spider’s centroid. Meanwhile, vertical coordinates were normalized by subtracting their mean value over time.

After centering joint coordinates, we then applied the Morlet continuous wavelet transform^39^ to capture spider movements.

The wavelet transforms described by Berman et. al.^39^ was defined as follows:

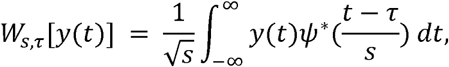

with

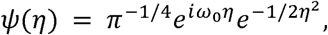

where *y*(*t*) is spider’s postural time series, *ω*_0_ = 5 is a non-dimensional parameter, *τ* is a point in time, and *s* is the time scale of interest as a function of frequency *f*:

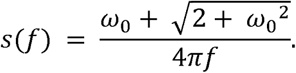

The power spectrum is:

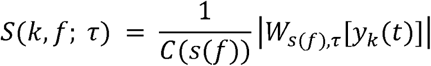

with the scalar function

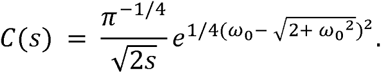

Finally, the frequency range used was between *f_min_* = 0.1 and the Nyquist frequency (*f_max_* = 50 Hz), with 50 frequencies space as follows:

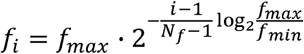

Wavelet analysis was applied on 20 joints in 4 legs for each side camera recordings. The wavelet spectrum was therefore had a dimension of 20 joints × 50 frequencies × 2 coordinates (vertical and horizontal movements) for each video.

### Stereotyped prey capture behavior motifs

We observed strong correlations among the wavelet representations of all four legs, as well as significant dependence between vertical and horizontal components. Additionally, low-frequency wavelets were primarily associated with noise, such as air current perturbations on the web. To reduce dataset complexity, we focused our analysis on five joint wavelets in the vertical coordinate from the spider’s left anterior leg, using 25 frequencies ranging from 2.38 to 50 Hz. This resulted in a wavelet spectrum of dimensions 5 joints × 25 frequencies × 1 coordinate × 21 recordings.

To identify stereotyped movements during prey-capture, we applied a Uniform Manifold Approximation and Projection (UMAP)^27^ for dimensionality reduction to five components, using 100 neighbors. From the wavelet spectrum, we observed 3 different motifs: no movement (static), middle frequency movement (2.38-10 Hz; crouching), and high frequency movement (10-50 Hz). Therefore, - means clustering^28^ with 3 clusters was applied on UMAP to identify 3 different behavioral motifs.

### Unsupervised Hidden Markov Model

To characterize spider prey-capture dynamics, we constructed a Hidden Markov Model with three hidden states, initializing the emission probabilities as multivariate Gaussian distributions based on the three motifs identified by K-means clustering. The transition matrix was randomly initialized. The model was then trained using the Baum-Welch algorithm, with a minimum of 50 and a maximum of 500 iterations. Note that true state labels were never used to train HMM. Therefore, the model is unsupervised.

### GLM-HMM

The GLM–HMM framework was originally proposed by Calhoun et al.^31^. We introduced two key simplifications: First, the filters were estimated using a linear behavior-triggered average. Second, the HMM transition probabilities were fixed so the GLM component was fully independent of the HMM.

The inputs to the GLM were two sensory signals from *Drosophila*: the normalized short-time Fourier transform (STFT) of fly, and the fly’s vertical velocity. The STFT power was Z-scored to account for variation in lighting angles across flies, which can influence light intensity and STFT power. The vertical velocity signal was low-pass filtered using a 25 Hz Butterworth filter to reduce high-frequency noise.

Instead of using an expectation-maximization algorithm to estimate sensory filters, we adapted the spike-triggered average (STA) from the linear-nonlinear (LN) model to a behavior-triggered average (BTA), which provides a more simple, computationally efficient, and non-parametric method to estimate stimulus-response mappings for each state. The spider’s behavioral states were predicted using a joint wavelet Hidden Markov Model (HMM). The BTA was defined as the average of sensory inputs during the 100-milisecond window preceding each behavioral state. Then, we took inner product of raw sensory inputs with linear filters from BTA and fit a logistic regression to estimate probability of each state.

Finally, we simply took the state with largest probability from the GLM as the input to HMM. The HMM structure was the same as what we describe in the previous section. We estimated the initial emission probability based on GLM prediction and randomly initialized the transition probability. The model was then trained using the Baum-Welch algorithm, with a minimum of 50 and a maximum of 500 iterations.

## Acknowledgements

We thank C. Li and E. Lin for helpful discussions and feedback. We are also grateful to A. Rabinovich, S. Gafrey, and T. Kolawole for their assistance in annotating spider joints. A.G. discloses support for the research of this work from NIH [R35GM124883], and NSF [2310707]. H.H. discloses support from The Ministry of Education (MOE) Taiwan Scholarship Program.

## Author Contributions

H.H. and A.G. conceived and designed the research strategy. A.C. designed the experimental apparatus and wrote the web-annotation software. H.H. built the experimental apparatus and optimized the web- annotation software. H.H. performed all experiments. H.H. and A.G. analyzed the data and prepared the manuscript.

## Data Availability

All software written for this study can be found at https://github.com/GordusLab/spider_prey_capture_paper.

## Reference

1. Jakobsen, L., Olsen, M. N. & Surlykke, A. Dynamics of the echolocation beam during prey pursuit in aerial hawking bats. Proc Natl Acad Sci U S A 112, 8118–8123 (2015).

2. Patterson, B. W., Abraham, A. O., MacIver, M. A. & McLean, D. L. Visually guided gradation of prey capture movements in larval zebrafish. J Exp Biol 216, 3071 (2013).

3. Roemschied, F. A. et al. Flexible circuit mechanisms for context-dependent song sequencing. Nature 622, 794–801 (2023).

4. Hindmarsh Sten, T., Li, R., Otopalik, A. & Ruta, V. Sexual arousal gates visual processing during Drosophila courtship. Nature 595, 549–553 (2021).

5. Schroeder, C. E., Wilson, D. A., Radman, T., Scharfman, H. & Lakatos, P. Dynamics of Active Sensing and perceptual selection. Curr Opin Neurobiol 20, 172–176 (2010).

6. Nelson, M. E. & MacIver, M. A. Sensory acquisition in active sensing systems. J Comp Physiol A Neuroethol Sens Neural Behav Physiol 192, 573–586 (2006).

7. Cellini, B. & Mongeau, J. M. Active vision shapes and coordinates flight motor responses in flies. Proc Natl Acad Sci U S A 117, 23085–23095 (2020).

8. Rucci, M., Iovin, R., Poletti, M. & Santini, F. Miniature eye movements enhance fine spatial detail. Nature 447, 851–854 (2007).

9. Johnson, M., Madsen, P. T., Zimmer, W. M. X., Aguilar De Soto, N. & Tyack, P. L. Beaked whales echolocate on prey. Proceedings of the Royal Society B: Biological Sciences 271, 383–386 (2004).

10. Amichai, E., Blumrosen, G. & Yovel, Y. Calling louder and longer: How bats use biosonar under severe acoustic interference from other bats. Proceedings of the Royal Society B: Biological Sciences 282, (2015).

11. Schnitzler, H. U., Moss, C. F. & Denzinger, A. From spatial orientation to food acquisition in echolocating bats. Trends Ecol Evol 18, 386–394 (2003).

12. Wachowiak, M. All in a Sniff: Olfaction as a Model for Active Sensing. Neuron 71, 962–973 (2011).

13. Stamper, S. A., Roth, E., Cowan, N. J. & Fortune, E. S. Active sensing via movement shapes spatiotemporal patterns of sensory feedback. Journal of Experimental Biology 215, 1567–1574 (2012).

14. Von der Emde, G. & Schwarz, S. Imaging of Objects through active electrolocation in Gnathonemus petersii. Journal of Physiology-Paris 96, 431–444 (2002).

15. Deora, T., Ahmed, M. A., Daniel, T. L. & Brunton, B. W. Tactile active sensing in an insect plant pollinator. Journal of Experimental Biology 224, (2021).

16. Prescott, T. J., Diamond, M. E. & Wing, A. M. Active touch sensing. Philosophical Transactions of the Royal Society B: Biological Sciences 366, 2989–2995 (2011).

17. Staudacher, E. M., Gebhardt, M. & Dürr, V. Antennal Movements and Mechanoreception: Neurobiology of Active Tactile Sensors. Advances in Insect Physiology vol. 32 (2005).

18. Arkley, K., Grant, R. A., Mitchinson, B. & Prescott, T. J. Strategy change in vibrissal active sensing during rat locomotion. Current Biology 24, 1507–1512 (2014).

19. Possomato-Vieira, José S. and Khalil, R. A. K. & Modeling, O. 2. 0: E. S. E. and S. Whisking mechanics and active sensing. Physiol Behav 176, 139–148 (2017).

20. Mortimer, B. A spider’s vibration landscape: Adaptations to promote vibrational information transfer in orb webs. Integr Comp Biol 59, 1636–1645 (2019).

21. Barth, F. G. & Geethabali. Spider vibration receptors: Threshold curves of individual slits in the metatarsal lyriform organ. Journal of Comparative Physiology IZ A 148, 175–185 (1982).

22. Klärner, D. & Barth, F. G. Vibratory signals and prey capture in orb-weaving spiders (Zygiella x-notata, Nephila clavipes; Araneidae). Journal of Comparative Physiology IZ A 148, 445–455 (1982).

23. Corver, A. Sensorimotor dynamics of the web-making behavior of the spider Uloborus diversus. Preprint at https://jscholarship.library.jhu.edu/handle/1774.2/69441 (2024).

24. Markow, T. A. & O’Grady, P. M. Drosophila Biology in the Genomic Age. Genetics 177, 1269–1276 (2007).

25. Mathis, A. et al. DeepLabCut: markerless pose estimation of user-defined body parts with deep learning. Nat Neurosci 21, 1281–1289 (2018).

26. Berman, G. J., Choi, D. M., Bialek, W. & Shaevitz, J. W. Mapping the stereotyped behaviour of freely moving fruit flies. J R Soc Interface 11, (2014).

27. McInnes, L., Healy, J. & Melville, J. UMAP: Uniform Manifold Approximation and Projection for Dimension Reduction. (2018).

28. Hartigan, J. A. & Wong, M. A. Algorithm AS 136: A K-Means Clustering Algorithm. Appl Stat 28, 100 (1979).

29. Corver, A., Wilkerson, N., Miller, J. & Gordus, A. Distinct movement patterns generate stages of spider web building. Current Biology 31, 4983–4997.e5 (2021).

30. Mearns, D. S., Donovan, J. C., Fernandes, A. M., Semmelhack, J. L. & Baier, H. Deconstructing Hunting Behavior Reveals a Tightly Coupled Stimulus-Response Loop. Current Biology 30, 54–69.e9 (2020).

31. Calhoun, A. J., Pillow, J. W. & Murthy, M. Unsupervised identification of the internal states that shape natural behavior. Nat Neurosci 22, 2040–2049 (2019).

32. Masters, W. M. Vibrations in the orbwebs of Nuctenea sclopetaria (Araneidae) - I. Transmission through the web. Behav Ecol Sociobiol 15, 207–215 (1984).

33. Barth, F. G. A Spider’s World: Senses and Behavior. Springer, Berlin (2002).

34. Landolfa, M. A. & Barth, F. G. Vibrations in the orb web of the spider Nephila clavipes: Cues for discrimination and orientation. J Comp Physiol A 179, 493–508 (1996).

35. Jones, T. K., Allen, K. M. & Moss, C. F. Communication with self, friends and foes in active-sensing animals. Journal of Experimental Biology 224, (2021).

36. Beleyur, T. & Goerlitz, H. R. Modeling active sensing reveals echo detection even in large groups of bats. Proc Natl Acad Sci U S A 116, 26662–26668 (2019).

37. Mitchinson, B. et al. Active vibrissal sensing in rodents and marsupials. Philosophical Transactions of the Royal Society B: Biological Sciences 366, 3037–3048 (2011).

38. Wallach, A. & Sawtell, N. B. An internal model for canceling self-generated sensory input in freely behaving electric fish. Neuron 111, 2570–2582.e5 (2023).

39. Berman, G. J., Choi, D. M., Bialek, W. & Shaevitz, J. W. Mapping the stereotyped behaviour of freely moving fruit flies. J R Soc Interface 11, (2014).

