## Supplementary figures and images for "Unsupervised discovery and predictive sensorimotor transformation of spider prey capture through active vibration sensing"

### Supplementary Figure 1

#### Supplementary Figure 1

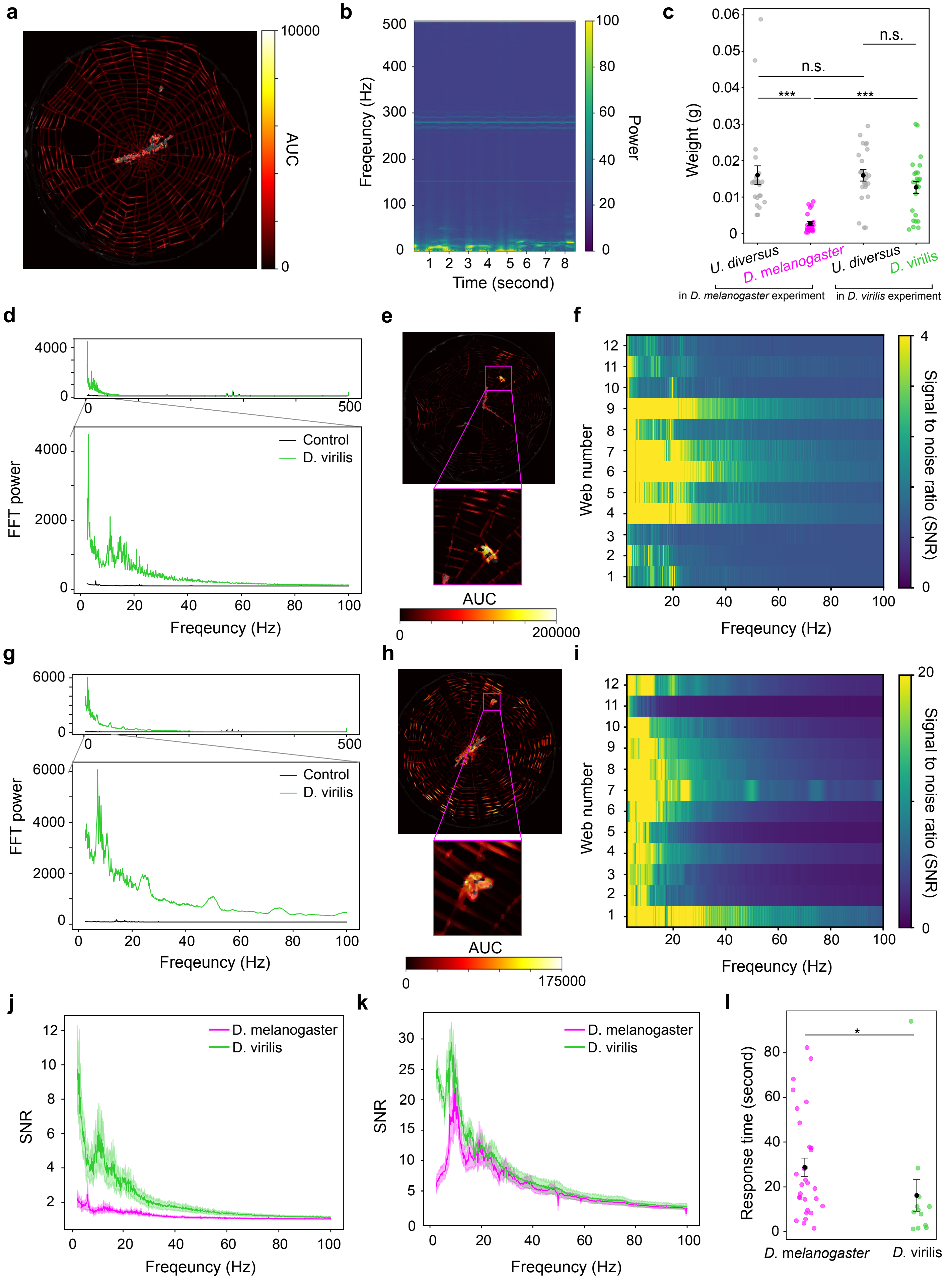

### Supplementary Figure 2

**Supplementary Figure 2**

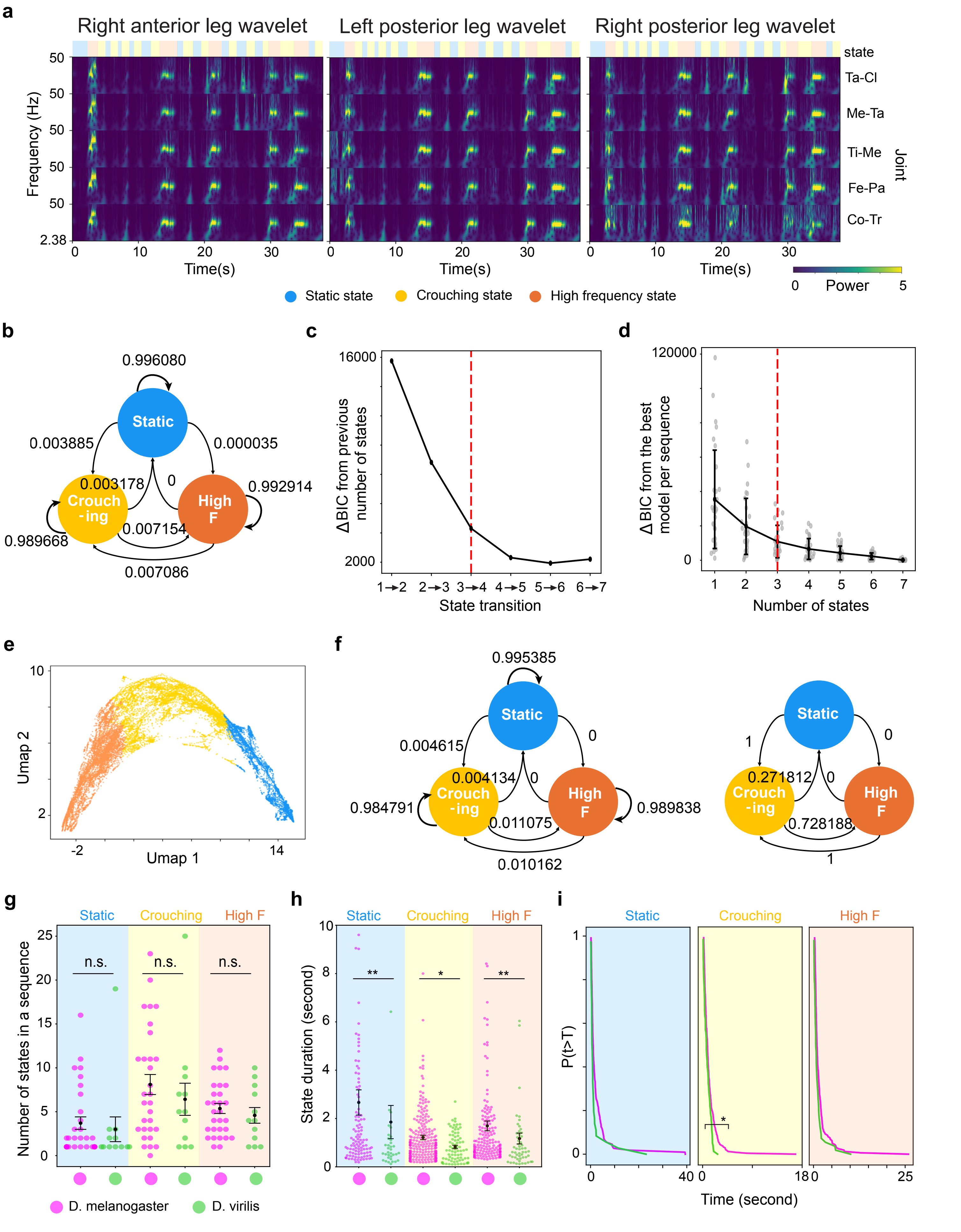

### Supplementary Figure 3

#### Supplementary Figure 3

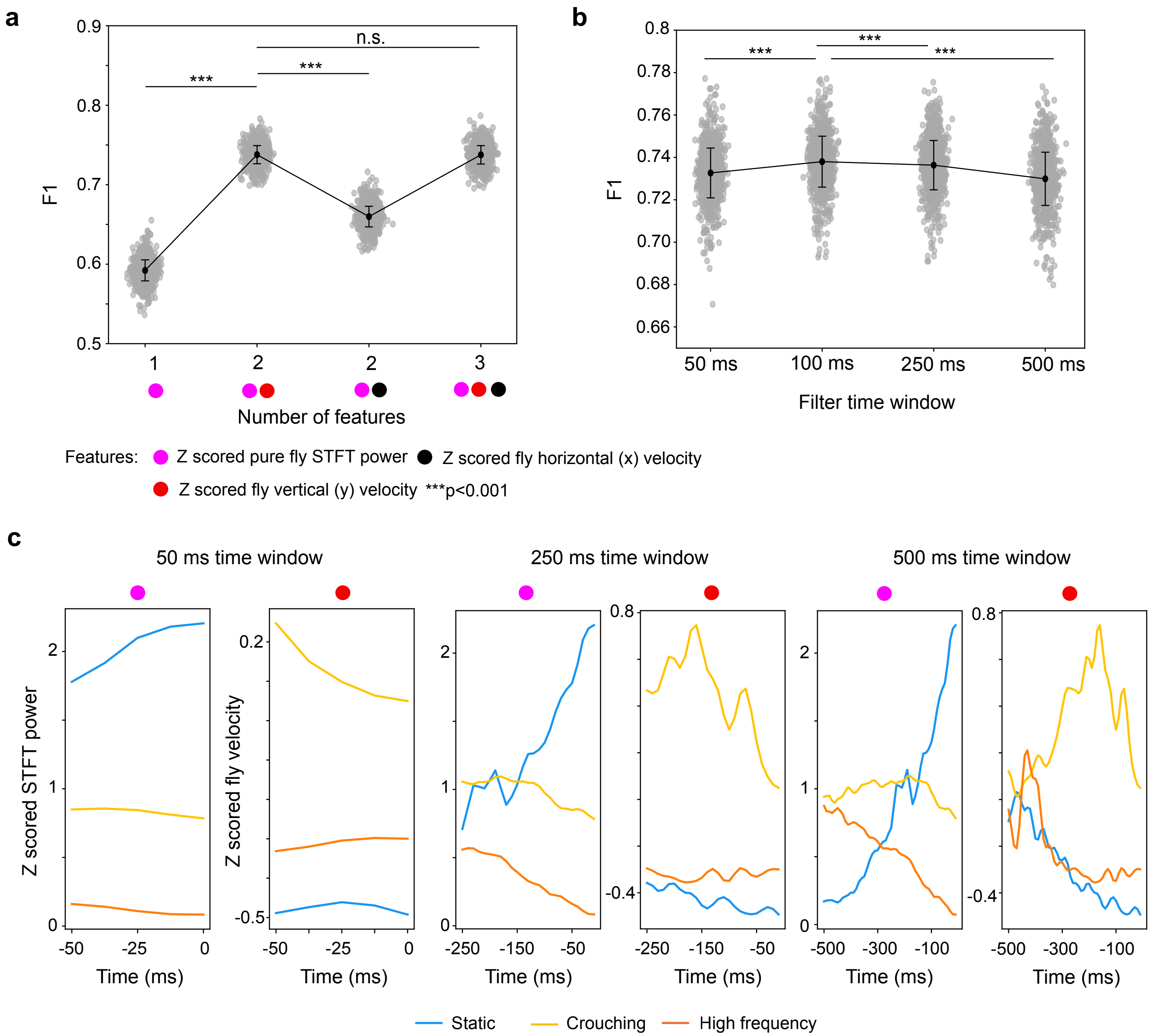
